# Predictive power of different *Akkermansia* phylogroups in clinical response to PD-1 blockade against non-small cell lung cancer

**DOI:** 10.1101/2024.08.21.608814

**Authors:** Peixin Fan, Mi Ni, Yu Fan, Magdalena Ksiezarek, Gang Fang

**Affiliations:** Department of Genetics and Genomic Sciences, Icahn School of Medicine at Mount Sinai, New York, NY, USA

**Keywords:** Non-small cell lung cancer (NSCLC), immunotherapy, *Akkermansia muciniphila*, phylogroup

## Abstract

Immune checkpoint blockade has emerged as a promising form of cancer therapy. However, only some patients respond to checkpoint inhibitors, while a significant proportion of patients do not, calling for the discovery of reliable biomarkers. Recent studies reported the importance of the gut microbiome in the clinical response to PD-1 blockade against non-small cell lung cancer (NSCLC), highlighting *Akkermansia muciniphila* as a candidate biomarker. Motivated by the genomic and phenotypic differences across *Akkermansia muciniphila* strains and *Akkermansia* (Akk) phylogroups (AmIa, AmIb, AmII, AmIII and AmIV), we analyzed fecal metagenomic sequencing data from four publicly available NSCLC cohorts (n = 575). Encouragingly, we found that patients’ responses to PD-1 blockade are significantly different across different Akk phylogroups, highlighting the relatively stronger association between AmIa and positive responses than the other phylogroups. Importantly, we also highlight the importance of considering the clinical heterogeneity among independent cohorts in across validation analysis. We built a machine learning model based on Akk gene profiles, which shed light on a group of Akk genes that may be associated with the response to PD-1 blockade. In summary, our study underlines the benefits of high-resolution analysis of Akk genomes in the search for biomarkers that may improve the prediction of patients’ responses to cancer immunotherapy.

## Introduction

Lung cancer is the third most common type of cancer in the US and the leading cause of cancer-related mortality worldwide.^1,2^ Non-small cell lung cancer (NSCLC) is the most prevalent form of lung cancer (84%).^3^ The emergence of immunotherapy, especially the immune checkpoint blockade, has revolutionized the cancer treatments with great promise, especially for NSCLC.^4,5,6^. However, only some patients experience durable responses to immune checkpoint inhibitors (ICIs), while others do not respond. For example, although NSCLC has a relatively higher response rate to ICIs than other cancer types, only ∼20% of NSCLC patients respond favorably to ICIs as monotherapy.^7^ Therefore, it is urgent to understand critical factors that influence the response to immunotherapy and discover predictive biomarkers that may guide personalized medicine.

Several promising biomarkers have been discovered for predicting responses to immunotherapy and have been incorporated into clinical practice, including the expression of immune checkpoint proteins (e.g., PD-1 or PD-L1), pre-existing immune responses, tumor mutation burden, and tumor-associated proteins.^7,8,9,10,11^ From a recent study, bronchoalveolar lavage fluid analysis of cellular profiles or secreted molecules can help identify patients with NSCLC who respond to ICIs.^12^ In addition, recent studies reported an association between patients’ gut microbiome and their responses to immunotherapies.^13,14^ Animal model-based studies found that antitumor activity of ICIs was lower in microbiome-depleted mice such as germ-free mice and mice treated with antibiotics, compared to specific pathogen-free mice.^15,16^ Patient cohort studies showed significant differences in gut microbiome composition between NSCLC, renal cell carcinoma, and melanoma patients who responded to anti-PD-1 therapy and those who did not.^16,17,18,19^ Fecal microbiota transplantation (FMT) was shown to influence the ICIs treatment of refractory melanoma and colorectal cancer in clinical trials,^20,21^ suggesting some specific gut microbes may serve as non-invasive predictive biomarkers.

For NSCLC, recent studies reported that *Akkermansia muciniphila*, a mucin-degrading gut commensal bacterium, was associated with better response to PD-1-based immunotherapy, and that it can enhance ICIs treatment in mice.^22,23,24^ Functionally, *A. muciniphila* is associated with beneficial effects on host metabolism and homeostatic immunity.^25,26^ It can also directly trigger intestinal adaptive immune responses through inducing T cell-dependent IgG1 and IgA.^24^ Immunogenic activity of lipid (a diacyl phosphatidylethanolamine with two branched chains) and peptides (e.g., Amuc_1100 and P9) produced by *A. muciniphila* were observed *in vitro.*^27,28^ Notably, several studies have demonstrated the genomic and phenotypic differences across *A. muciniphila* strains, which initiated further classification into *Akkermansia* (Akk) phylogroups (primarily AmIa, AmIb, AmII, AmIII and AmIV). Among these phylogroups, only AmIa and AmIb represent *A. muciniphila,* while the remaining are being actively characterized as potential new *Akkermansia* species.^29,30,31,32^ Different Akk phylogroups were shown to have significant phenotypic differences that are associated with different host-microbe interactions: growth rates in mucin, binding to epithelial surfaces, and activation of innate immune responses.^29,30,31,32,33^. A recent study also identified correlations between *Akkermansia* species and subspecies and human health outcomes.^32^ Consistently, two different Akk strains showed different capacities to modulate differentiation of regulatory T cells (Tregs) in mice.^16^ Based on these genetic and phenotypic variations among Akk strains, we hypothesized that NSCLC patients carrying different phylogroups of Akk in their gut microbiome have different responses to immunotherapy.

To test this hypothesis, we first determined the distribution of Akk phylogroups in NSCLC metagenomics datasets of 575 samples from four publicly available cohorts^16,23,34^ and evaluated the predictive power of different Akk phylogroups for patients’ responses to PD-1 blockade.

Furthermore, we performed a rigorous analysis and highlighted a subset of genes in the Akk pangenome that are enriched in patient responders.

## Results

### Akk phylogroups across four cohorts of NSCLC patients with ICI treatment

We performed a meta-analysis across four publicly available NSCLC cohorts (*C1*: *Routy* et al. Discovery, n = 60; *C2*: *Routy* et al. Validation, n = 27; *C3: Derosa* et al. 2022, n = 338, and *C4: Derosa* et al.2024, n = 150) that used fecal shotgun metagenomics to study the associations between Akk and response to PD-1 blockade. All raw shotgun metagenomic sequencing data were processed consistently using the same bioinformatics pipeline (STAR Methods). The available metadata of the four cohorts was summarized in Table S1.

We first re-evaluated the associations between the presence/absence of Akk genus (Akk^+^ or Akk^-^) or major *A. muciniphila* species-level genome bins (SGBs) (SGB9226^+^ or SBG9226^-^) determined by MetaPhlAn 4^35^, STAR Methods) and response to PD-1 blockade against NSCLC. The number of clean sequencing reads per sample did not have a significant difference between Akk^+^ and Akk^-^ samples (*P*_C1_ = 0.55, *P*_C2_ = 0.73, *P*_C3_ = 0.11, *P*_C4_ = 0.13, non-parametric Wilcoxon rank-sum test, Fig. S1A-D) or between SGB9226^+^ and SBG9226^-^ samples (*P*_C1_ = 0.13, *P*_C2_ = 0.32, *P*_C3_ = 0.65, *P*_C4_ = 0.22, non-parametric Wilcoxon rank-sum test, Fig. S1E-H) across the four cohorts. Consistent with previous studies, the Akk^+^ and SGB9226^+^ groups had higher (compared to the Akk^-^ and SBG9226^-^) proportion of progression-free patients [complete response (CR), partial response (PR), or stable disease (SD), Fig. S2A-D] and objective response (CR and PR, Fig. S2E-H) in all cohorts. Noteworthy, in the *C2* cohort, all the 5 patients with CR or PR were characterized as Akk^+^ group (*P* = 0.016, Fig. S2F). Besides, the variation in response rate between Akk^-^ and Akk^+^ groups was larger than that between SGB9226^-^ and SBG9226^+^ groups, suggesting other Akk bins may be associated with patients’ response to PD-1 blockade. We also revaluated the consistency of the associations between the relative abundance of Akk and SGB9226 (instead of the presence/absence of Akk and SGB9226) and patients’ response, across the four cohorts, as *Derosa* et al. 2022 reported that among Akk (SGB9226) positive patients, those with high abundance of Akk (SGB9226) (>4.799%) showed significantly lower response rate compared to patients with low abundance of Akk (SGB9226).^23^ However, this phenomenon was only observed in C2 and C3 cohorts, while Akk^high^ and SGB9226^high^ patients showed much higher response rate than Akk^low^ and SGB9226^low^ patients in both C1 (Akk^high^ vs Akk^low^: 80% vs 55.56%, SGB9226^high^ vs SGB9226^low^: 75% vs 62.5%) and C4 cohorts (Akk^high^ vs Akk^low^: 83.33% vs 55.71%, SGB9226^high^ vs SGB9226^low^: 84.62% vs 54.55%) (Fig. S3), indicating that high relative abundance of Akk or SGB9226 was not consistently associated with negative response to PD-1 blockage. These analyses supported the use of the presence/absence of Akk in the following analyses, which is consistent with a recent study that reported bacterial presence/absence is more robust than bacterial relative abundance in microbiome-based association studies.^36^ Overall, NSCLC patients carrying Akk and SGB9226 (presence vs. absence) had a higher response rate to PD-1 blockade, consistently across the four cohorts.

To distinguish Akk phylogroups in the meta-analysis, we first built a comprehensive Akk pangenome including diverse Akk strains (216 Akk isolate genomes from the NCBI database) that include five known Akk phylogroups (Fig. S4A; STAR Methods). Out of the 220 Akk^+^ samples determined by MetaPhlan version 4 across the four cohorts, 24 (out of 32), 8 (out of 12),122 (out of 178), and 66 (out of 88) samples were assigned with an Akk phylogroup by PanPhlan 3 (STAR Methods),^37^ respectively (Fig. 1A-B, Fig. S4B-D, C1: 10 AmIa, 10 AmIb, and 4 AmII; C2: 4 AmIa, 3 AmIb, and 1 AmII; C3: 58 AmIa, 37 AmIb, 21 AmII, and 6 AmIV; C4: 25 AmIa, 26 AmIb, 14 AmII, and 1 AmIV). 90 samples are Akk^+^ according to MetaPhlAn 4 but failed to be assigned to the Akk phylogroups, and all of them had low Akk abundance (0.096% ± 0.01%). In addition, the previous work showed there is usually only a single phylogroup in an individual’s gut microbiome, and the coexistence of multiple Akk phylogroups is very rare.^30^ Therefore, the reason that certain Akk^+^ samples were not assigned to a specific Akk phylogroup was mainly due to the low Akk abundance but not a mixture of different phylogroups. Though MetaPhlAn 4 can effectively differentiate between AmI (*A. muciniphila* SGB9226), AmII (*A. muciniphila* SGB9228) and AmIV (Akkermansia sp. KLE1798 SGB9224), it cannot differentiate between AmIa and AmIb, the two major phylogroups of Akk, which are both assigned to SGB9226 by MetaPhlan 4. Consistent across the four cohorts, AmIa and AmIb are the two most prevalent Akk phylogroups, representing typical *A. muciniphila* species, the major *A. muciniphila* bin SGB9226 and the most common Akk species found in the human microbiome^26^, followed by AmII (SGB9228) and AmIV (SGB9224). Therefore, Akk phylogroups determined by PanPhlan 3 was used to evaluate their effects on response to PD-1 blockade in this study.

**Figure 1.**
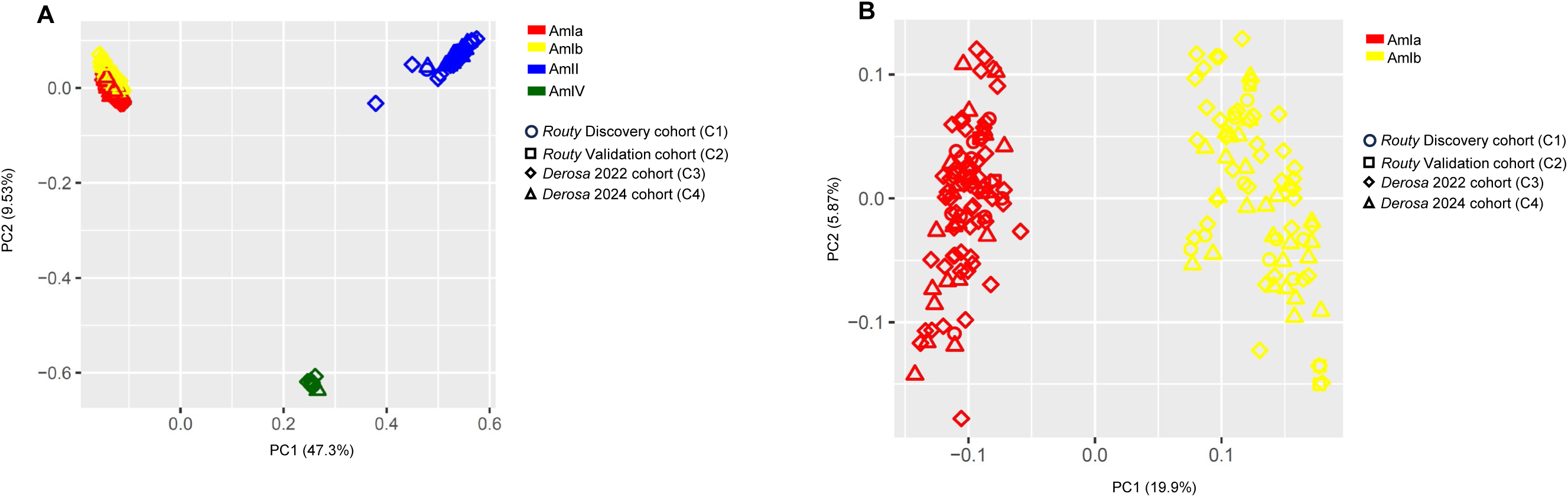
Distinguishing NSCLC patients carrying different phylogroup of Akk. **(A)** The Akk phylogroups carried by NSCLC patients from four cohorts. **(B)** The AmIa and AmIb phylogroup carried by NSCLC patients from four cohorts. The four cohorts are *Routy* Discovery cohort (n = 60), *Routy* Validation cohort (n = 27), *Derosa* 2022 cohort (n = 338), and *Derosa* 2024 cohort (n = 150). Principal component analysis (PCA) was performed based on the *Akkermansia* gene presence/absence matrix from PanPhlan, and the results are projected in the plot.

### Akk phylogroups have different predictive power for patient response to PD-1 blockade

We compared patients’ response rate between NSCLC patients carrying different Akk phylogroups using a logistic regression model (with covariates; STAR Methods) and found that AmIa is associated with better response to PD-1 blockade across all the four NSCLC cohorts and that higher proportion of responders than non-responder was observed in patients carrying AmIa (Fig. 2). Specifically, the rates of responders (CR, PR or SD) in the AmIa group were 80% in *C1* (*P*_AmIa_ = 0.037, Fig. 2A), 75% in *C2* (*P*_AmIa_ = 0.075, Fig. 2B), 58.62% in *C3* (*P*_AmIa_ = 0.016, Fig. 2C), and 56% in C4 (*P*_AmIa_ = 0.607, Fig. 2D), respectively. However, this trend differed in patients carrying AmIb and AmII across cohorts. Although patients with AmIV also showed a higher proportion of responders, they represented only a small subset of patients and were found exclusively in the C3 and C4 cohorts. Besides, a higher proportion of responders were observed in patients carrying AmIa compared to those carrying AmIb in C1-C3 cohorts but not in the C4 cohort. Notably, line of therapy (LIGNE) showed a significant effect on response to PD-1 blockade in the C4 cohort (*P* = 0.012) that a greater proportion of patients with LIGNE =1 was responder compared to patients with LIGNE > 1 (Fig. S5C-D), which was not observed in C3 cohort (*P* = 0.72, Fig. S5A-B). Different from patients with LIGNE > 1 in C4 cohort and other three cohorts, among patients with LIGNE = 1 in C4 cohort, the association between presence of Akk or SGB9226 and better response was not observed (Fig. S5C-D). However, among patients with LIGNE > 1 in C4, the better response in Akk^+^ patients (compared to Akk^-^ patients), SGB9226^+^ patients (compared to SGB9226^-^ patients) (Fig. S5D), and patients carrying AmIa (compared to AmIb) were detected (Fig. S6B). Moreover, the better response of patients carrying AmIa compared to patients carrying AmIb was more apparent among patients with LIGNE > 1 compared to patients with LIGNE =1 in C3 cohort as well (Fig. S6A), suggesting an interaction between LIGNE and Akk phylogroups with respect to patients’ response to PD-1 blockade.

**Figure 2.**
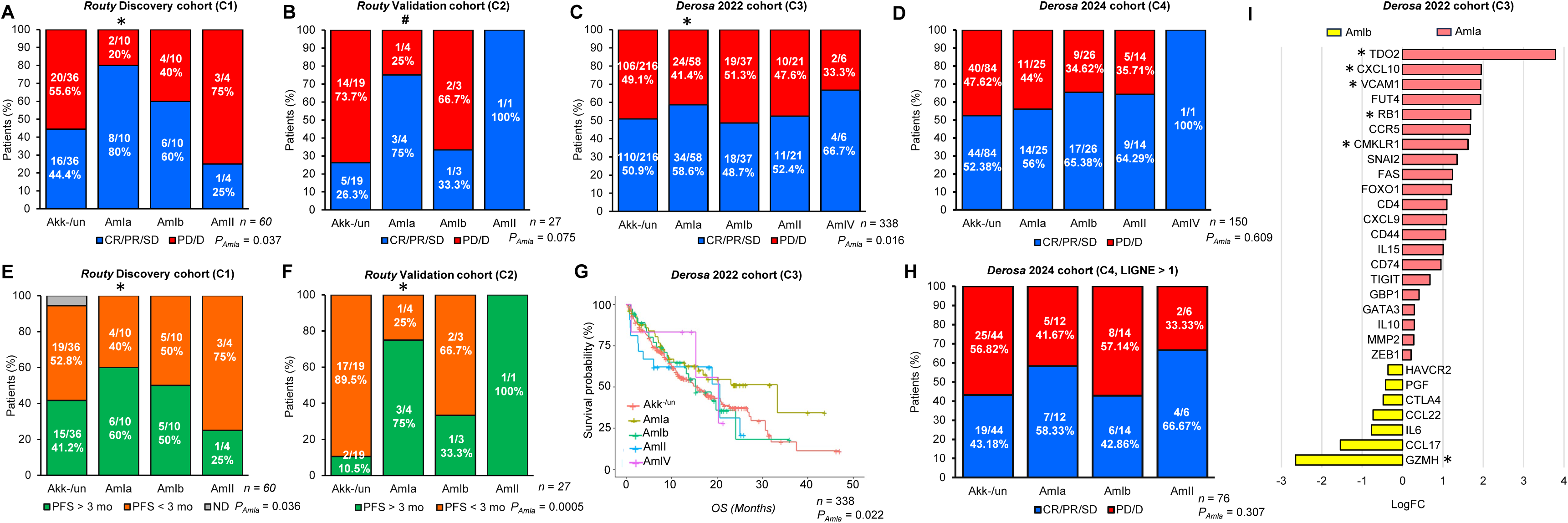
NSCLC patients carrying different phylogroup of Akk varied in response to ICI treatment. (A to. **D)** Correlations between phylogroup of Akk in the stool of NSCLC patients and response to PD-1 blockade. AmIa group had higher proportion of patients being CR, PR, or SD in *Routy* Discovery cohort (n = 60, *P* = 0.037) (**A**), *Routy* Validation cohort (n = 27, *P* = 0.075) (**B**), and *Derosa* 2022 cohort (n = 338, *P* = 0.016) (**C**) and Derosa 2024 cohort (n = 150, *P* = 0.609) (**D**). In *Routy* Validation cohort, AmIa group showed significantly higher proportion of patients being CR or PR (*P* = 9.04×10^-8^). CR; complete response. PR; partial response. SD; stable disease. PD; progressive disease. D; death. Data were analyzed using generalized linear model. (**E**-**G)** Correlations between phylogroup of Akk in stool of NSCLC patients and survival length. AmIa group had higher proportion of patients which had progression-free survival (PFS) longer than 3 months in *Routy* Discovery cohort (n = 60, *P* = 0.036) (**E**) and *Routy* Validation cohort (n = 27, *P* = 0.0005) (**F**), and had longer overall survival in *2022 Derosa* cohort (n = 338, *P* = 0.022) (**G**). The response rate in LIGNE > 1 patients carrying different Akk phylogroups in 2024 *Derosa* cohort (C4) (**H**). Data were analyzed using generalized linear model or Cox proportional-hazards regression. (**I**), Bar graph showing most immune-related genes that were expressed at higher level in lung tumor of patients carrying Akk had higher expression level in AmIa group. Differential expression analysis was conducted using voom method. Akk-/un represents for the samples that are either detected as Akk negative using MetaPhlAn 4 or detected as Akk positive in MetaPhIAN 4 but not in PanPhlan. *, *P* < 0.05; ^#^, 0.05 *< P* < 0.1.

We also evaluated the logistic regression models that included presence/absence of Akk phylogroup determined by PanPhlan vs. presence/absence of Akk bins determined by MetaPhlan to predict objective response (CR and PR) using corrected Akaike information criterion (AICc) ^38^ which measures the relative prediction error between regression models (the smaller value the better model). Consistently across the four cohorts, the logistic regression models that included Akk phylogroup have better accuracy and AICc than the models that used Akk species: C1: AICc = 64.246 (Akk phylogroup) vs 71.740 (Akk SGB); *C2*: AICc = 13.097 (Akk phylogroup) vs 34.877 (Akk SGB); *C3*: AICc = 253.745 (Akk phylogroup) vs 259.286 (Akk SGB); *C4*: AICc = 106.978 (Akk phylogroup) vs 107.380 (Akk SGB). We further compared patient survival across different Akk phylogroups. In the *C1* cohort, 60% of patients carrying AmIa had progression-free survival (PFS) longer than 3 months (vs. 43.75% of the other patients, *P*_AmIa_ = 0.036, Fig. 2E). In the *C2* cohort, 75% of patients carrying AmIa had PFS longer than 3 months (vs. 33.33% of AmIb patients, *P*_AmIa_ = 0.0005, Fig. 2F). Consistently, in the C3 cohort, AmIa patients had longer overall survival (OS) than patients not carrying AmIa based on Cox proportional-Hazards regression analysis (*P*_AmIa_ = 0.022, Fig. 2G). It is worth noting that both AmIa and AmIb are prevalent in the three NSCLC cohorts, yet AmIa is associated with better patients’ response and survival than AmIb, consistently across the C1-C3 cohorts, and C4 cohort patients with LIGNE > 1 (Fig. 2H). We further investigated relative abundance of different Akk phylogroups and their influence on response to PD-1 blockade across the four cohorts.

There was no significant difference in relative abundance between Akk phylogroups across all the cohorts (*P_C1_* = 0.32, *P_C2_*= 0.23, *P_C3_* = 0.55, *P_C4_* = 0.89, Fig. S7). The phenomenon of better response in Akk^low^ patients was observed in patients carrying different Akk phylogroups in C3 cohort and patients carrying AmIb or AmII in C2 cohort, but not in C1, C4, and patients carrying AmIa in C2 cohort (Fig. S8), suggesting presence/absence of Akk phylogroup rather than relative abundance is more consistently associated with response to PD-1 blockade. We also performed a meta-analysis across the four cohorts (C1, C2, C3 excluding Akk^high^ patients, and C4 LIGNE > 1 patients) and observed that the presence of AmIa showed a positive trend toward association with treatment response (*P* = 0.096), with a 95% confidence interval of (-0.21, 2.56) and moderate heterogeneity (Tau² = 0.45) after adjusting for other covariates.

*Derosa* et al. reported in the *C3* cohort that tumor microenvironment differed between Akk^+^ patients and Akk^-^ patients, highlighting several differentially expressed genes.^23^ To evaluate if Akk phylogroups and patients’ differential response and survival rate correlate with patients’ tumor microenvironment, we examined the gene expression levels of 28 immune-related genes between patients with AmIa and AmIb phylogroups. Interestingly, 21 of the 28 genes (*P_binomial_* = 0.006) that were expressed higher in Akk^+^ patients than Akk^-^ patients showed greater expression level in patients carrying AmIa compared to those in AmIb, with 5 of them showing significant difference (*P_adj_* < 0.05, Fig. 2I). The *TDO2* gene, which is the interferon fingerprint and associated with checkpoint pathway, showed the largest difference between AmIa and AmIb group (LogFC = 3.79, *P_adj_* = 0.008). *CXCL10* (LogFC = 1.95, *P_adj_* = 0.049), related to type II interferon signaling, *VCAM1* (LogFC = 1.94, *P_adj_* = 0.015), related to leukocyte migration, *RB1* (LogFC = 1.69, *P_adj_* = 0.033) related to tumor marker, and *CMKLR1* (LogFC = 1.62, *P_adj_* = 0.048), related to dendritic cell and macrophage also had higher expression in AmIa than AmIb. These analyses suggest that the tumor microenvironment genes associated with Akk^+^ patients have more pronounced changes in their expression level in the patients carrying the AmIa phylogroup than the AmIb, consistent with their stronger anti-tumor immune response to ICI.

Taken together, these analyses suggest that the presence of AmIa phylogroup in NSCLC patients’ gut microbiome is associated with better responses than the other Akk phylogroups to PD-1 blockage, survival and tumor microenvironment.

### Akk gene-based prediction of response to anti-PD1 blockade

Building on the observations above that NSCLC patients carrying AmIa were associated with better response than the other phylogroups, we further hypothesized that AmIa genomes may be enriched for certain genes that are positively associated with the response to ICI. It is also worth noting that not all the patients carrying AmIa responded to PD-1 blockade while some patients with other Akk phylogroups did respond to PD-1 blockade. This heterogeneity within a phylogroup is not unexpected considering the genomic differences across strains within each phylogroup. Therefore, we further performed an Akk gene-level analysis to better understand the different predictive power between Akk phylogroups for patients’ response to PD-1 blockage.

To investigate whether Akk genes enhance predictive power of response to PD-1 blockage, we built a machine-learning model (model A) with patients’ Akk phylogroup, patients’ metadata (covariates: age, gender, antibiotics, number of clean reads, which are available in all the four cohorts) to predict response (CR/PR/SD or PD/D). We compared the accuracy of model A with another model (model B) built with the addition of presence or absence of Akk genes. We first trained the two models using all the available Akk^+^ samples with defined phylogroups in the combined *C1*, *C2*, and *C3* cohort (154 Akk^+^ samples), and tested the two models on the C4 cohort (66 Akk^+^ samples). The test accuracy of the model B is 0.57, which is higher than the accuracy of model A without gene-level data (0.52) (Fig. S9). Because the response rate was also influenced by Akk abundance in the C3 cohort and by LIGNE in the C4 cohort, we aimed to reduce heterogeneity across cohorts. To achieve this, we trained two additional models using all available Akk-positive samples from the combined C1, C2, and C3 cohorts after removing Akk-high samples (n = 121). Model 1 excluded all Akk-related genes, whereas Model 2 retained the Akk genes. We then tested both models on the C4 cohort after excluding patients with LIGNE = 1 (n = 42 Akk-positive samples). The model 1 using random forest had an area under the receiver operating characteristic (AUROC) value of 0.57 (Fig. 3A), while the model 2 had an AUROC of 0.67 (Fig. 3A). The test accuracy of the model 2 is 0.64, which is higher than the accuracy of model 1 without gene-level data (0.52). The improvements in LOOCV accuracy of the training model with the addition of Akk gene feature indicate the association between certain Akk genes and patients’ response to PD-1 blockade can provide better predictive power than Akk phylogroups itself.

**Figure 3.**
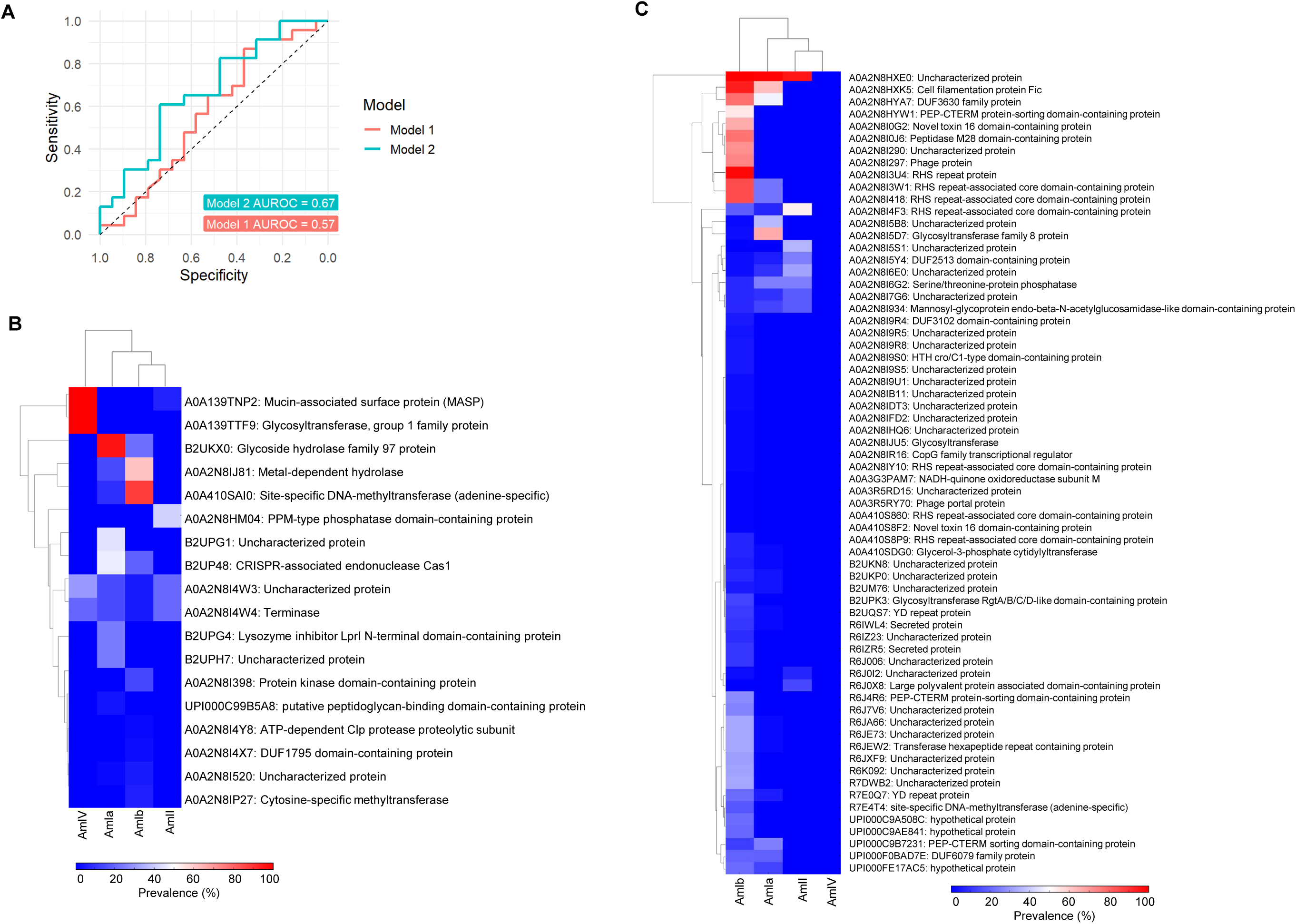
Akk gene-based model improved performance to predict response to PD-1 blockade and potential Akk genes that are associated with response to PD-1 blockade. **(A)** The area under the receiver operating characteristic (AUROC) of the random forest models trained to predict response using Akk phylogroup, patients’ metadata, number of clean reads, and without (model 1) or with (model 2) presence/absence of Akk genes in the combined C1 (n = 24), C2 cohort (n = 8), and C3 cohort excluding Akk^high^ (n = 89), and tested on C4 cohort excluding LIGNE =1 patients (n = 42). **(B)** The heatmaps showing the distribution of responder-associated Akk genes across Akk phylogroups. Only Akk genes that significantly or tended to be enriched in responders (*P*_Chi-square_ < 0.10) and with variable importance score (VIC) higher than 1 in the random forest model (model 2) were included in the heatmap. **(C)** The heatmaps showing the distribution of non-responder-associated Akk genes across Akk phylogroups. Only Akk genes that significantly or tended to be enriched in non-responders (*P*_Chi-square_ < 0.10) and with variable importance score (VIC) higher than 1 in the random forest model (model 2) were included in the heatmap.

To investigate which genes contribute significantly to improving the predictive power and their distribution across different phylogroups, we combined the 163 Akk^+^ patients with defined Akk phylogroup across the four cohorts (excluding the Akk^high^ patients in C3 and patients with LIGNE =1 in C4 cohorts) and constructed a new model for the ranking of important Akk genes. This model is based on patients’ metadata, number of clean reads, Akk phylogroups, and presence and absence matrix of all the 3263 pan-Akk genes, selected using strict criteria to avoid ambiguous mapping of genes from other species to Akk (STAR Methods). Among Akk phylogroups, only presence/absence of AmIa showed a variable importance score (VIS) greater than 0 (0.11), indicating its greater predictive importance compared to other phylogroups. There were 84 important Akk genes (VIS > 1) showing significantly different or tendency in prevalence between responders and non-responders (*P_chi-square_* < 0.10, Table S2), with 18 enriched in the responders (Fig. 3B) and 66 enriched in the non-responders (Fig. 3C), respectively. Among the 18 Akk genes enriched in the responders, 7 genes had highest prevalence in AmIb, followed by 6 genes in AmIa, 3 genes in AmIV, and 2 gene in AmII (Fig. 3B). These responder-associated genes included several carbohydrate-active enzymes (CAZy) encoding genes, such as genes encoding glycoside hydrolase family 97 protein, which had the highest VIS (10.21, *P_chi-square_* =0.03) and is present in 96.30% of AmIa reference strains, 21.56% of AmIb strains, but not in AmII and AmIV strains; glycosyltransferase, group 1 family protein, which is uniquely present in AmIV reference strains (100%). Moreover, two immune-related Akk genes including mucin-associated surface protein (MASP) present in all AmIV strains and few AmII strains (6.70%), and lysozyme inhibitor LprI N-terminal domain-containing protein present in a portion of AmIa strains (24.07%) only, were also enriched in responders.

For the 66 Akk genes enriched in non-responders over responders, 54 genes had highest prevalence in AmIb, followed by 9 genes in AmII and 3 genes in AmIa (Fig. 3C). These genes encode several bacteriophage related proteins, such as phage portal protein and phage protein, as well as bacterial toxin-related proteins, such as Novel toxin 16 domain-containing protein, multiple RHS repeat proteins that play important roles in bacterial virulence, such as RHS repeat protein and RHS repeat-associated core domain-containing protein, and secreted proteins, which all had the highest prevalence in AmIb. Among them, the gene encodes secreted protein (UniRef90_R6IWL4) had the highest VIS (100, *P_chi-square_* = 0.00017). Besides, two genes encode YD repeat protein, which are associated with toxin delivery were much more prevalent in AmII (UniRef_R7E0Q7, 34.48%) and AmIa (UniRef_B2UQS7, 64.48%), respectively. Moreover, nonresponder-associated Akk genes also include certain CAZy encoding gene, particularly those encode glycosyltransferase, glycosyltransferase family 8 protein, and Glycosyltransferase RgtA/B/C/D-like domain-containing protein, which were also more prevalent in AmIb compared to the other Akk phylogroups.

These results suggest a link between carbohydrate utilization of gut *Akkermansia* and response to immunotherapy of hosts, likely reflecting variation in intestinal mucosal environments, particularly glycoside diversity, between responders and nonresponders, which probably results from their distinct immune and metabolic status, that favor growth of different Akk phylogroups.^39,40^ The patients dominated by AmIa and AmIV phylogroup may also benefit from bacterial glycoside fermentation end products, such as short-chain fatty acids, that enhance their response to PD-1 blockade.^41,42^. However, patients dominated by AmIb, particularly the AmIb strains with virulence factors could be a reflection of their immune suppression which enabled the proliferation of potential harmful Akk strains which in turn further disrupted their immunity. AmIa contained the second most genes enriched in responders and second fewest genes enriched in nonresponders, while AmIb contained much more genes enriched in nonresponders, which is consistent with the observation that patients with AmIa were more likely than AmIb to be responders. Although few patients carried the AmIV strain, it harbors several unique responder-associated genes and lacks nonresponder-associated genes, which likely explains why its presence corresponds to the highest proportion of responders in the C3 and C4 cohorts.

To summarize, we found that the model that incorporate Akk genes has enhanced performance in the prediction of patients’ response to PD-1 blockade. Importantly, the gene-level association analysis (with patient response and with Akk phylogroups) helped elucidate the difference in the predictive power of different Akk phylogroups in clinical response to PD-1 blockade against NSCLC.

## Discussion

In this study, we assessed the difference between Akk phylogroups with respect to their predictive capacity in responses to PD-1 blockade in NSCLC patients. Through the delineation of Akk phylogroups and an extensive meta-analysis across four previously published cohorts, we unveiled diverse Akk phylogroups among NSCLC’s patients, highlighting the better power of the AmIa phylogroup, consistently across four independent cohorts, in predicting responses to immunotherapy compared to the other phylogroups. Also consistently, we observed stronger association between AmIa and patient survival and the RNA expression of patients’ genes relevant to tumor microenvironment. Using a machine learning model predicting patients’ response based on Akk gene profiles, we searched for the genes associated with AmIa’s superior predictive capabilities, and highlighted a number of genes predominantly prevalent in responders and nonresponders to PD-1 blockade and their varied distribution across Akk phylogroups. AmI harbored more genes associated with responder compared to AmII and AmIV, but within AmI, AmIb harbored more genes associated with nonresponders than AmIa, which is consistent with the observation that patients carrying AmIa are more likely to be responders compared to patients carrying AmIb. These revelations underscore the importance of discerning Akk phylogroups among the major *Akkermansia muciniphila* bin and Akk genes in disease association analysis, which may help the development of innovative microbiome-centric diagnostic tools and personalized medicine approaches for PD-1 blockage in NSCLC patients.

Our integration and analysis of the four cohorts also emphasized some limitations and the need for further validations through larger, independent studies, considering the complexity of NSCLC and patients’ response to PD-1 blockage. First, although our analytical models used three co-variants shared across the three cohorts (age, gender, antibiotics), we think the collection and incorporation of additional covariates is paramount in these analyses, given their potential to introduce confounding effects. For instance, cohorts C1 and C2 predominantly comprised patients who underwent chemotherapy before PD-1 blockade, whereas cohort C3 had a higher proportion of patients treated with radiotherapy before PD-1 blockade. These variations might explain the relatively better predictive power of AmIa in C1 and C2 than in C3, even though the three cohorts consistently highlight the importance of AmIa. Second, while this study was focused on the microbiome, we also recognize the substantial importance of host factors such as ECOG performance status, PD-L1 expression, LIGNE, tumor gene expression data, in predicting patients’ response to PD-1 blockage.^23^ Although ECOG performance status, PD-L1 expression, and LIGNE are available in the C3 and C4 cohorts, they were not part of the C1 and C2 cohorts in our study. Given the complexity of NSCLC and patients’ response to PD-1 blockage, future development in this field necessitates studies with a holistic approach, integrating both microbiome and host gene expression data to decipher their relationships and their joint predictive power^24,43,44^. Besides, due to the limitation of annotated Akk gene based on our current knowledge, the functions of response-associated sequence clusters need to be further explored.

While Akk has persistently demonstrated a link to responses in NSCLC patients undergoing PD-1 blockade across varied cohorts, several other bacterial species like *Eubacterium hallii*, *Bifidobacterium adolescentis*, and *Parabacteroides distasonis* have also been reported, albeit with less consistency across cohorts^23,45,46^. This discrepancy might arise from the possibly more subtle yet legitimate associations these bacteria have with PD-1 blockade responses or due to the inherent patient heterogeneity observed across diverse cohorts. Indeed, the complexity of responses to PD-1 blockade in NSCLC is multifaceted, interweaving numerous factors from both the human host and the gut microbiome. While Akk is a significant player in inducing intestinal adaptive immune responses during homeostasis, it’s plausible that other bacterial species also play integral roles in this biological puzzle. With this rationale, we made an attempt to assess the predictive capabilities of additional bacterial species, probing their association with responses to PD-1 blockade (Supplementary Text).

*Monoglobus pectinilyticus* exhibited a consistent positive association with responders across C1-C3 NSCLC cohorts (Supplementary Text and Fig. S10). Besides, using metagenomic data of all the patients in the four cohorts, we attempted to identify overlap gene categories shared by Akk and other bacterial species associated with response to PD-1 blockade. However, we did not identify any gene categories that showed consistent associations with response to PD-1 blockade across. This suggests that the *Akkermansia*-associated genes linked to response may not be functionally replaceable by homologous genes from other species occupying the same ecological niche. Future work is imperative to validate this finding on large and independent cohorts as well as the complementarity between Akk and additional bacterial species.

Beyond its association with responses to PD-1 blockade, Akk has also been implicated in a spectrum of other diseases, including obesity, type 2 diabetes, and inflammatory bowel disease, among others.^47,48,49^ The insights from this study also suggest the use of similar analytical strategies to investigate the predictive capabilities of distinct Akk phylogroups in relation to these other diseases and their respective treatment outcomes. For example, the better response to PD-1 blockade of renal cell carcinoma (RCC) patients carrying AmIa than other phylogroups (Fig. S11).^16^ Given the recognition of Akk as a promising candidate in the development of next-generation probiotics, it is important to attain a nuanced understanding of the various Akk phylogroups, their specific genes, and genomic variations^50^ across a range of diseases, which could unravel transformative insights and facilitate the advancement of innovative therapeutic interventions and proactive preventive measures.

## STAR Methods

### Public populations of NSCLC patients

Raw gut metagenomic sequencing data of four populations (C1: *Routy* et al. Discovery, n = 60; C2: *Routy* et al. Validation, n = 27; C3: *Derosa* et al. 2022, n = 338, and C4: *Derosa* et al. 2024, n = 150) were downloaded from the Sequence Read Archive (SRA) under the BioProject accession PRJEB22863 (https://www.ncbi.nlm.nih.gov/bioproject/?term=PRJEB22863), PRJNA751792 (https://www.ncbi.nlm.nih.gov/bioproject/?term=PRJNA751792), and PRJNA1023797 (https://www.ncbi.nlm.nih.gov/bioproject/?term=PRJNA1023797) mainly from three recently published papers.^16,23,34^ The raw tumor RNA sequencing data were downloaded from the SRA under accession the NCBI accession GSE182328 (https://www.ncbi.nlm.nih.gov/geo/query/acc.cgi?acc=GSE182328) from *Derosa* et al. (2022).^23^

### Identification of Akk presence and Akk phylogroup in the stool metagenomes

The KneadData tool (v0.10.0) was used to perform quality control on raw metagenomic sequencing data. Briefly, adapters and low-quality reads were removed using Trimmomatic (v0.39), and repetitive sequences were trimmed using the TRF (v4.09.1). The human reads were detected by mapping remaining reads to the human genome (hg37) by bowtie2 (v.2.2.5) and removed. Presence and relative abundance of *Akkermansia* genus and *Akkermansia* species-level genome bins (SGBs) were determined and estimated by MetaPhlAn 4 (v4.0) with ChocoPhlAn database (vJun23_202307).^35^ PanPhlan pipeline (v3.0) was applied to profile *A. muciniphila* strains within stool metagenomes.^37^ A total of 216 available *A. muciniphila* isolate genomes (fasta file) were download from NCBI and were used to build a comprehensive pangenome with UniRef90 ID using PanPhlan pangenome exporter (https://github.com/SegataLab/PanPhlAn_pangenome_exporter). Due to high genetic diversity within *A. muciniphila* strains, this species has been suggested to be classified into four species-level phylogroups, and we considered the UniRef genes families-level pangenome as *Akkermansia* (Akk) pangenome. Then each metagenomic sample was mapped against the generated Akk custom pangenome (panphlan_map.py). The coverages of all gene positions were detected and extracted using samtools (v1.16), and then integrated to a gene family coverage profile for each sample. Pangenome-related Akk strains were predicted to exist if coverage curves display a satisfying plateau using the least stringent threshold of panphlan_profile.py (--min_coverage 1, --left_max 1.70, --right_min 0.30). Principal coordinates analysis (PCoA) of Jaccard distance was performed on UniRef90 ID presence and absence binary matrix of metagenomic samples using ggplot2 package in the R (v4.3) to distinguish phylogroup of Akk strains according to cluster with references.

### Evaluation of correlations between Akk presence/absence, Akk phylogroup and response to ICI treatment

Generalized linear model was applied to evaluate the effects of presence/absence or relative abundance of Akk genus, or Akk SGBs, or Akk phylogroups on the response. The patients were classified as responders including complete response (CR, Disappearance of all target lesions. Any pathological lymph nodes (whether target or non-target) must have reduction in short axis to <10 mm), partial response (PR, At least a 30% decrease in the sum of diameters of target lesions, taking as reference the baseline sum diameters), and stable disease (SD, Neither sufficient shrinkage to qualify for PR nor sufficient increase to qualify for progression disease, taking as reference the smallest sum diameters while on study) and non-responders including progressive disease (PD, At least a 20% increase in the sum of diameters of target lesions, taking as reference the smallest sum on study (this includes the baseline sum if that is the smallest on study). In addition to the relative increase of 20%, the sum must also demonstrate an absolute increase of at least 5 mm. the appearance of one or more new lesions is also considered progression) and death (D) using the best clinical response according to Response Evaluation Criteria in Solid Tumors version (RECIST) 1.1 according to the previous study.^16,51^ Akk presence or Akk phylogroup with sequencing depth (number of raw reads) and patients’ metadata (age, gender, time point, antibiotic usage, BMI, ECOG performance status, PD-L1 expression, LIGNE, smoking if available) were included as independent variables. Responders, objective response (CR and PR), or survival lengths (PFS > 3month in C1 and C2 cohorts) were served as dependent variables. Corrected Akaike information criterion (AICc) and accuracy were used to compare the quality of models. The survival analysis for C3 cohort was conducted using the Cox proportional-hazards regression.

### Tumor RNA sequencing data analysis

To investigate relationships between the expression of immune-related genes in tumors and Akk phylogroup the patients were carrying, the differences in Akk-associated immune-related genes reported by previous study was analyzed using the published tumor RNA sequencing data^23^. The low-quality reads and adapters were removed from raw RNA sequencing data using Trim Galore (v0.6.6). The remaining reads were mapped to human GRCh38 cDNA reference transcriptome using STAR (v2.7.2a). The number of reads mapping to each gene was counted using featureCounts (v2.0.1). The differentially expressed genes (DEGs) between patients carrying AmIa (n = 3) and AmIb (n = 5) were analyzed in R using voom package (v3.38.14).

### Construction and evaluation of the machine-learning model based on Akk genes profile

To determine whether Akk gene profiles could enhance the prediction of response to PD-1 blockade, the random forest models were built using R caret package (v 6.0-94). The patients from the four cohorts with Akk phylogroup assigned were included in this analysis. The training models were constructed with leave one out cross validation to distinguish responders and non-responders using the combined C1 (n = 24), C2 (n = 8), and C3 cohort excluding or not the Akk^high^ patients (n = 89 or n=122) as the training dataset and tested on the C4 cohort excluding or not LIGNE =1 patients (n = 42 or n = 66). The features used for model building consist of number of clean reads, patient metadata (age, gender, usage of antibiotics), and Akk profile (phylogroup, presence and absence of Akk only genes). The UniRef genes that only exist in *Akkermansia* were considered as Akk only genes. Briefly, the metadata for the full UniRef90 gene set was retrieved from the UniRef website (http://www.uniprot.org/uniref), which includes the organism/species information. All the Akk-related genes was extracted by using the Organism tag ("*Akkermansia*"). These Akk genes were classified into different categories based on the number of homologous. The genes that only existed in Akk were pulled out as Akk only genes. The models were optimized by tuning the mtry (number of randomly selected features to be sampled at each iteration) and the ntree (number of decision tree). The area under the receiver operating characteristic (AUROC) and accuracy were used to evaluate the performance of the training models.

### Identification of Akk genes associated with response to PD-1 blockade

To identify important Akk genes associated with response to PD-1 blockade, another random forest machine learning model was trained using the metadata and Akk profile of combined C1 (n = 24), C2 (n = 8), C3 cohort excluding the Akk^high^ patients (n = 89), and C4 cohort excluding LIGNE =1 patients (n = 42), with leave one out cross validation. The variable importance scores were extracted for each feature in the model, and distribution of critical Akk genes with high score (> 1) with significant difference or tendency present in responders and non-responders (*P_chi-square_*< 0.10) were determined across phylogroups. In addition, we developed a custom pipeline to test whether gut microbial genes with similar functions in non-Akkermansia species are also associated with ICI response. Gene abundances were estimated using HUMAnN v3.0.4 (http://huttenhower.sph.harvard.edu/humann) with the UniRef90 database (version 201901b, DIAMOND-annotated full reference). Per-sample gene tables were merged, and metagenomic association analysis was performed using MaAsLin2.^52^ Genes significantly associated with ICI response (*P* < 0.1) were mapped to UniRef50 clusters and grouped into functional categories based on gene or protein names. These were then compared to *Akkermansia*-linked genes through shared UniRef50 IDs. We retained genes showing consistent directions of association and overlapping or known functional categories. Functional enrichment was summarized at the category level, and statistical significance was assessed using a one-sided Mann–Whitney U test against randomly sampled background categories.

## Code Availability

The customized scripts of this study as an AkkPD1 package at GitHub (https://github.com/fanglab/AkkPD1) with detailed instructions and demo/test data.

## Acknowledgements

We thank Edward Mead, Yanchun Zhang and other members in the Fang Lab for their helpful feedback. This work was supported by R35 GM139655 (G.F.) from the National Institutes of Health. G.F. is a Hirschl Research Scholar by Irma T. Hirschl/Monique Weill-Caulier Trust, and a Nash Family Research Scholar. This work was also supported in part through the computational resources and staff expertise provided by the Department of Scientific Computing at the Icahn School of Medicine at Mount Sinai.

## Author Contributions

P.F. and M.Ni. performed all the data analyses. P.F. and G.F. designed the meta-analyses across the three cohorts. Y.F. and M.K. assisted in the Akk pangenome analysis. G.F. conceived and supervised the project. P.F. and G.F. wrote the manuscript with input and comments from all co-authors.

