## Supplementary information for "Predictive power of different *Akkermansia* phylogroups in clinical response to PD-1 blockade against non-small cell lung cancer"

**Other bacterial species complemented Akk to predict the response to PD-1 blockade**

Considering the complexity of gut microbiome and immunotherapy, patients’ response to PD-1 blockade may involve additional bacteria beyond Akk. With this rationale, we further evaluated the predictive power of other bacterial species and their complementarity with Akk in the association with patients’ response to PD-1 blockade. First, we used a generalized linear regression model to evaluate the effect of each individual bacterial species for their association with response to PD-1 blockade across the three cohorts (Fig. S4; Methods). The presence of *Monoglobus pectinilyticus* showed positive association with responders consistently in all the three NSCLC cohorts (*P_C1_=*0.090, *P_C2_*=0.089, *P_C3_*=0.00683; meta-analysis *P =* 0.0261). The presence of some additional bacteria showed positive association with responders (OR, PR or SD) in two of the three NSCLC cohorts: *Bifidobacterium bifidum (P_C2_=0.043, P_C3_=0.048)*, *Olsenella scatoligenes (P_C2_=0.027, P_C3_=0.009)*, and *Flavonifractor sp* An10 *(P_C2_=0.054, P_C3_=0.014)*. Second, we evaluated whether the combination of Akk and *Monoglobus pectinilyticus* will improve the accuracy of predicting response to PD-1 blockade. Specifically, we compared the model based on Akk phylogroup with or without the incorporation of *Monoglobus pectinilyticus* in the model. *M. pectinilyticus (Mp)* as an additional covariate to Akk phylogroup improved the model accuracy: AIC = 575.49 (Akk phylogroup) vs 568.79 (Akk phylogroup + Mp); Accuracy = 61.58% (Akk phylogroup) vs 63.87% (Akk phylogroup + Mp). It is worth noting that, although the improvement in predictive accuracy was very modest, the incorporation of *M. pectinilyticus* explained 16.1% additional more responders compared to AmIa, the Akk phylogroup positively enriched in responders (Fig. S4B). *M. pectinilyticus* has functional specialization for pectin degradation and utilization in human gut^1^, while its influence on human immunity is still largely unknown. These data suggest that in addition to Akk, other specific bacterial species may provide complementary predictive power as possible biomarkers for patients’ response to PD-1 blockage.

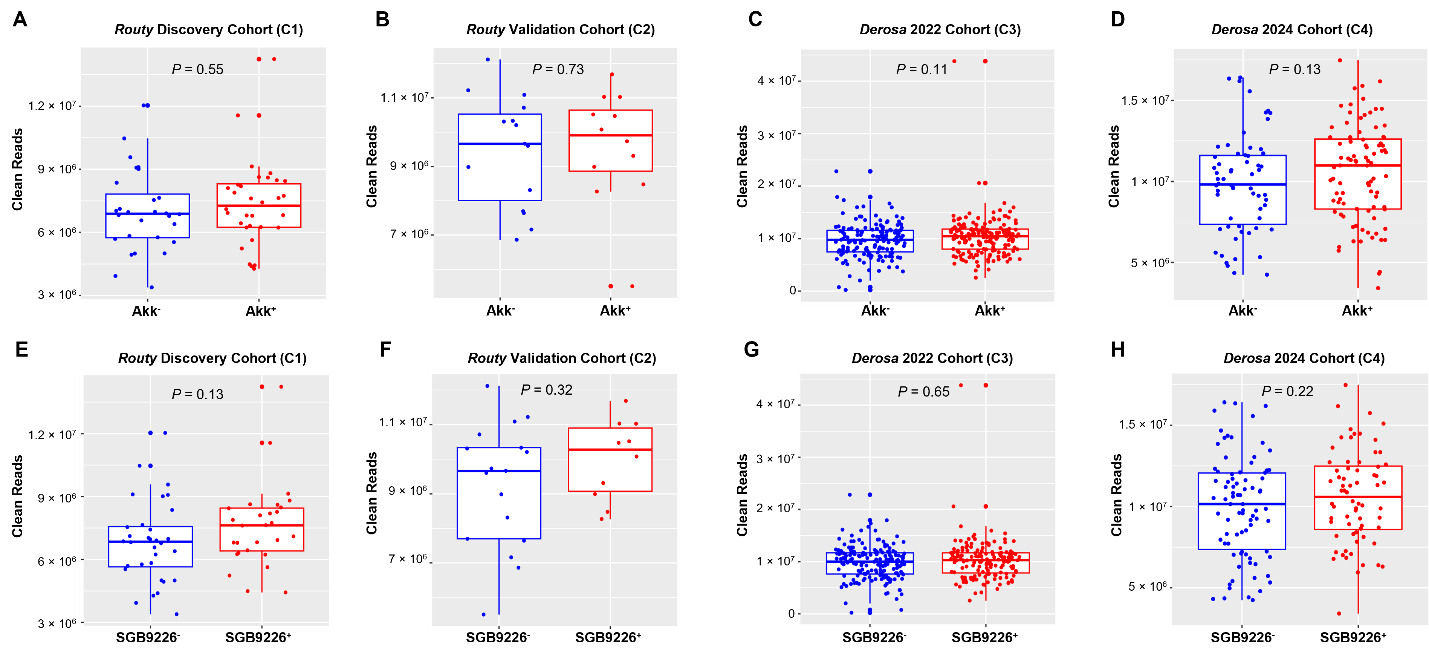

**Figure S1. The clean reads numbers between *Akkermansia* genus positive (Akk^+^) and negative (Akk-) samples or between major *Akkermansia muciniphila* bin (SGB9226^+^) positive and negative (SGB9226^-^) samples across four NSCLC cohorts.** The clean reads numbers between Akk^-^ and Akk^+^ samples in *Routy* Discovery cohort (C1) (**A**), *Routy* Validation cohort (C2) (**B**), *Derosa 2022* cohort (C3) (**C**), and *Derosa 2024* cohort (C4) (**D**). The clean reads numbers between SGB9226^-^ and SGB9226^+^ samples in *Routy* Discovery cohort (C1) (**E**), *Routy* Validation cohort (C2) (**F**), *Derosa* 2022 cohort (C3) (**G**), and *Derosa* 2024 cohort (C4) (**H**). The absence/presence of Akk or SGB9226 was analyzed using MetaPhlAn version 4. The statistical analysis was conducted using the non-parametric Wilcoxon rank-sum test.

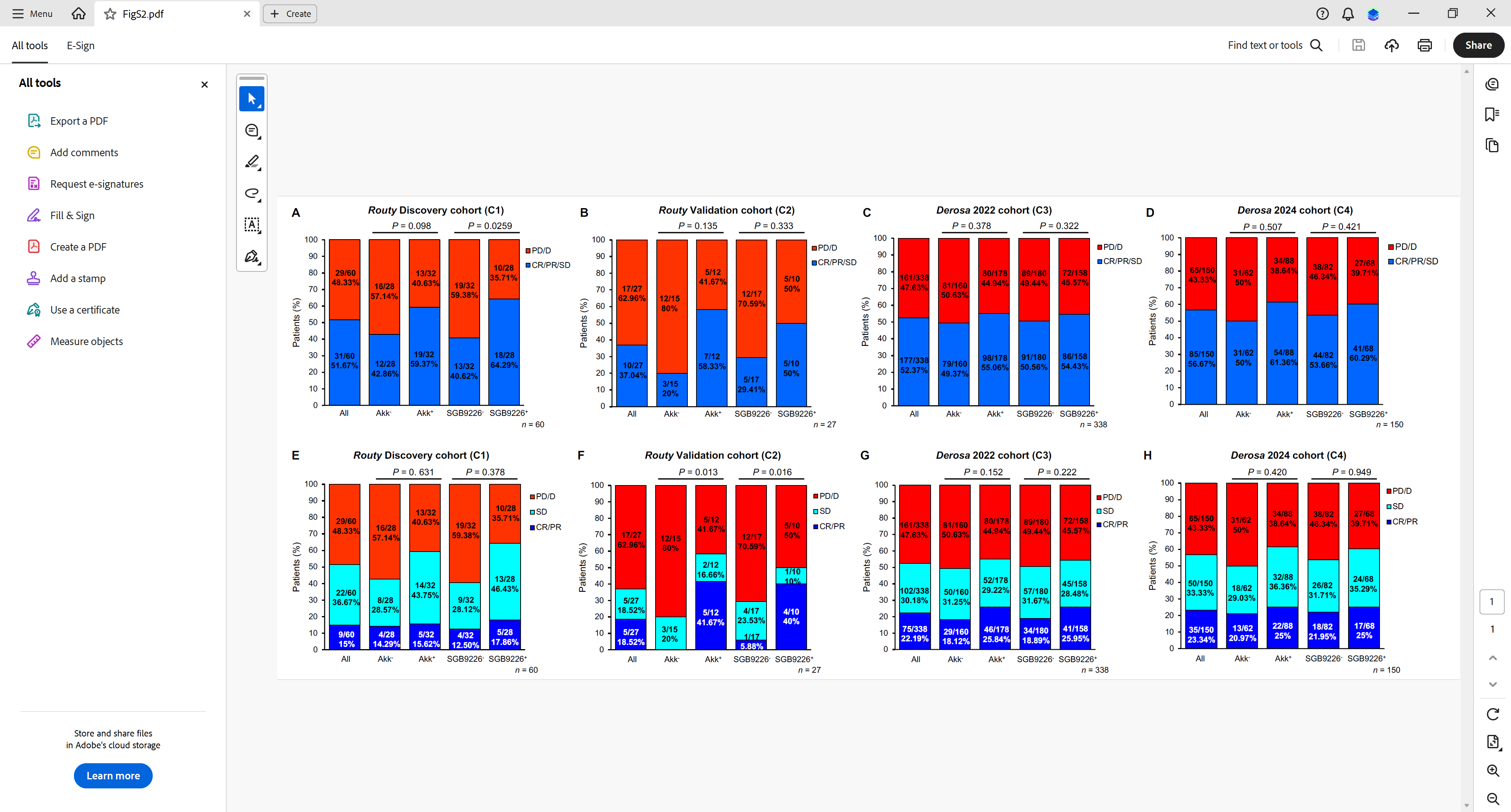

**Figure S2. Examination of the relationship between clinical response to PD-1 blockade against NSCLC and presence of *Akkermansia* genus and major *A. muciniphila* bin SGB9226 in gut of patients across four NSCLC cohorts.** Differences in proportion of responders between patients with or without carrying *Akkermansia* genus (Akk^+^ vs Akk^-^) or major *A. muciniphila* bin SGB9226 (SGB9226^+^ vs SGB9226^-^) in *Routy* Discovery cohort (C1) (**A**), Routy Validation cohort (C2) (**B**), *Derosa* 2022 cohort (C3) (**C**), and *Derosa* 2024 cohort (C4) (**D**). The responders include NSCLC patients showing complete response (CR), partial response (PD), or stable disease (SD) after PD-1 blockade. The nonresponders include NSCLC patients showing progressive disease (PD) or death (D). Differences in objective response rates (proportion of CR and PR) between patients that carry and not carry *Akkermansia* genus (Akk^+^ vs Akk^-^) and between patients that carry and not carry major *A. muciniphila* bin SGB9226 (SGB9226^+^ vs SGB9226^-^) in *Routy* Discovery cohort (C1) **(E)**, Routy Validation cohort (C2) (**F**), *Derosa* 2022 cohort (C3) (**G**), and *Derosa* 2024 cohort (C4) (**H**).

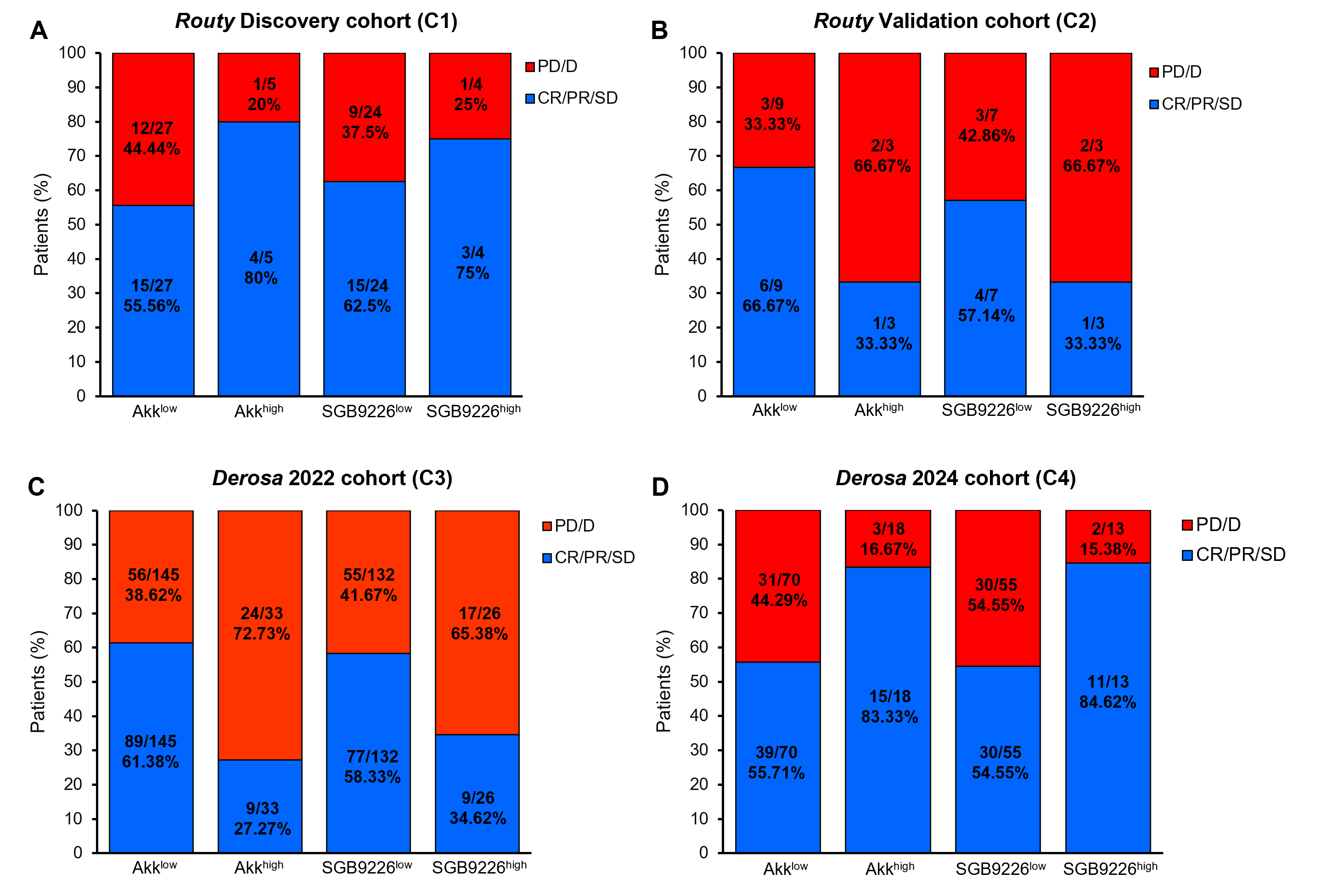

**Figure S3. Examination of the relationship between clinical response to PD-1 blockade against NSCLC and relative abundance *Akkermansia* genus and major *A. muciniphila* bin SGB9226 in gut of patients across four NSCLC cohorts.** Differences in proportion of responders between patients that carried high (> 4.799%) and low relative abundance (0 < Akk or SGB9226 < 4.799%) of *Akkermansia* genus (Akk^high^ vs Akk^low^) or major *A. muciniphila* bin SGB9226 (SGB9226^high^ and SGB9226^low^*)* in *Routy* Discovery cohort (C1) (**A**), Routy Validation cohort (C2) (**B**), *Derosa* 2022 cohort (C3) (**C**), and *Derosa* 2024 cohort (C4) (**D**). The responders include NSCLC patients showing complete response (CR), partial response (PD), or stable disease (SD) after PD-1 blockade. The nonresponders include NSCLC patients showing progressive disease (PD) or death (D).

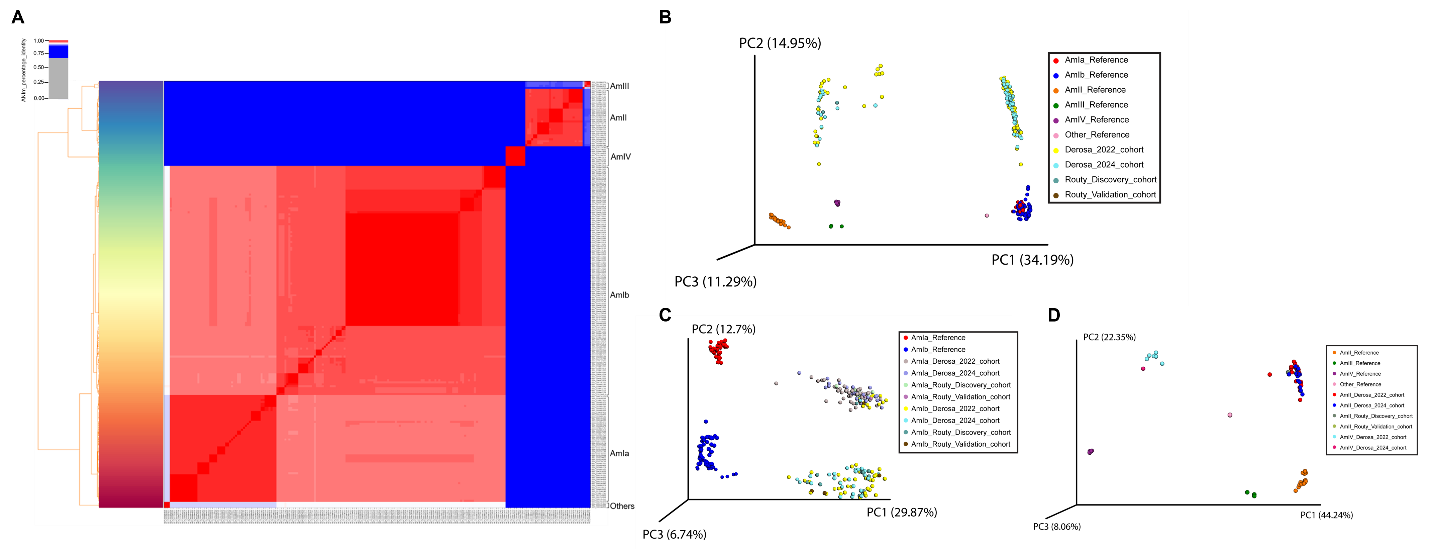

**Figure S4. The custom PanPhlAn pangenome built for *Akkermansia muciniphila* and assignment of *Akkermansia muciniphila* phylogroup for fecal metagenome across patients across four NSCLC cohorts.** (**A**) The heatmap showing the average nucleotide identity (ANI) percentage identity among 216 *Akkermansia muciniphila* genomes collected from NCBI database. These 216 *Akkermansia muciniphila* strains were classified into AmIa (n = 54), AmIb (n = 116), AmII (n = 29), AmIII (n = 4), AmIV (n = 10), and others (n = 3). (**B**) The principal coordinate analysis (PCoA) plot generated according to the presence/absence of UniRef90 gene families in fecal metagenomes from patients across four NSCLC cohorts against custom built *Akkermansia muciniphila* PanPhlAn pangenome. The patients were mainly separated into two clusters, one is close to AmIa and AmIb strains, and the other one close to AmII, AmIII, and AmIV. To further distinguish the *Akkermansia muciniphila* phylogroups, principal coordinate analysis (PCoA) was conducted for NSCLC patients that are in AmI cluster (**C**) and those are in AmII, AmIII, and AmIV cluster (**D**).

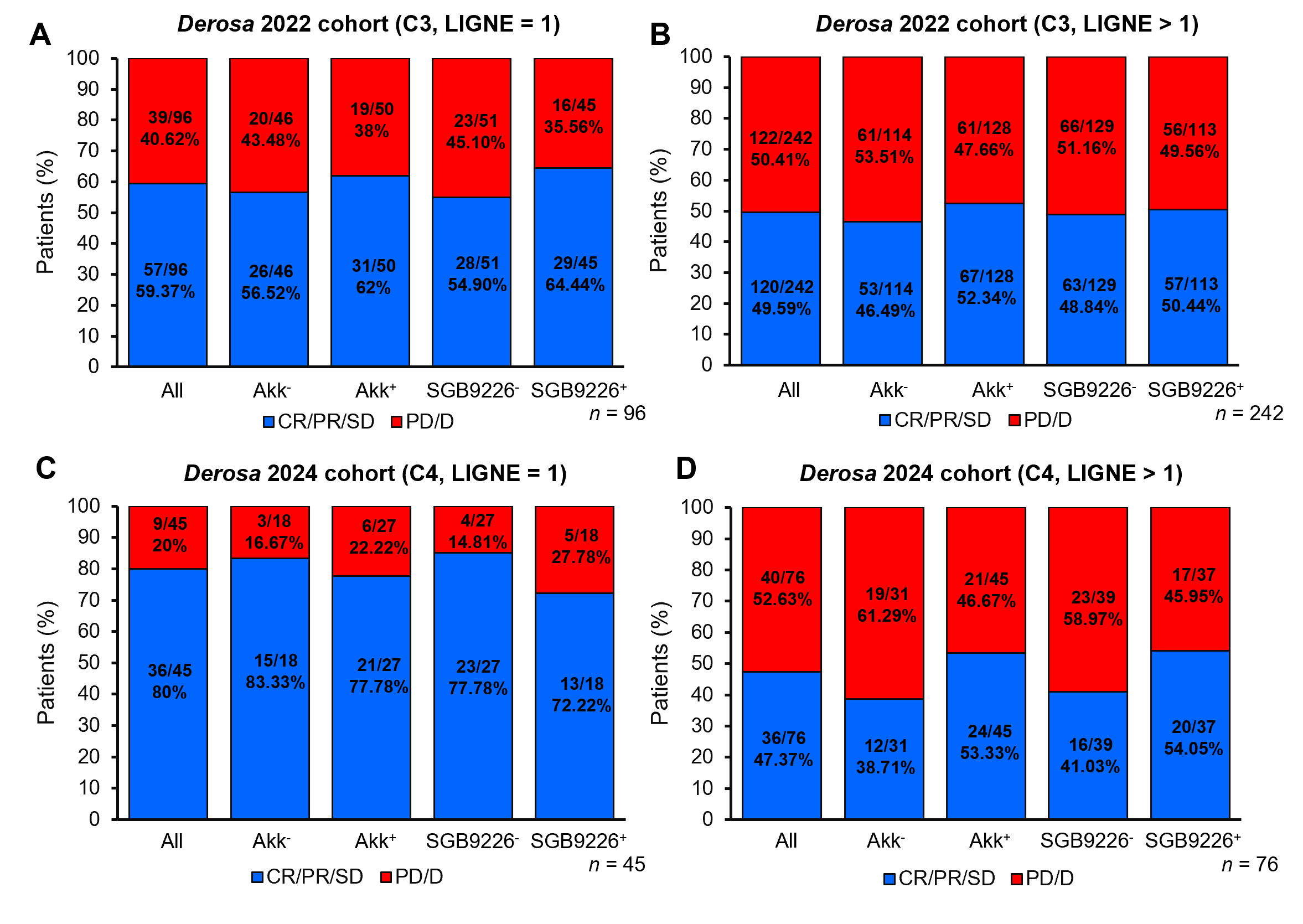

**Figure S5. The effect of LIGNE on the response to PD-1 blockade in patients with or without carrying *Akkermansia* genus, major *A. muciniphila* species-level genome bin (SGB9226) in two NSCLC cohorts.** Differences in response to PD-1 blockade between patients with and without carrying *Akkermansia* genus, SGB9226 among patients with LIGNE =1 (**A**) or LINGE > 1 (**B**) in *Derosa* 2022 cohort (C3). Differences in response to PD-1 blockade between patients with and without carrying *Akkermansia* genus, SGB9226 among patients with LIGNE =1 (**C**) or LINGE > 1 (**D**) in *Derosa* 2024 cohort (C4).

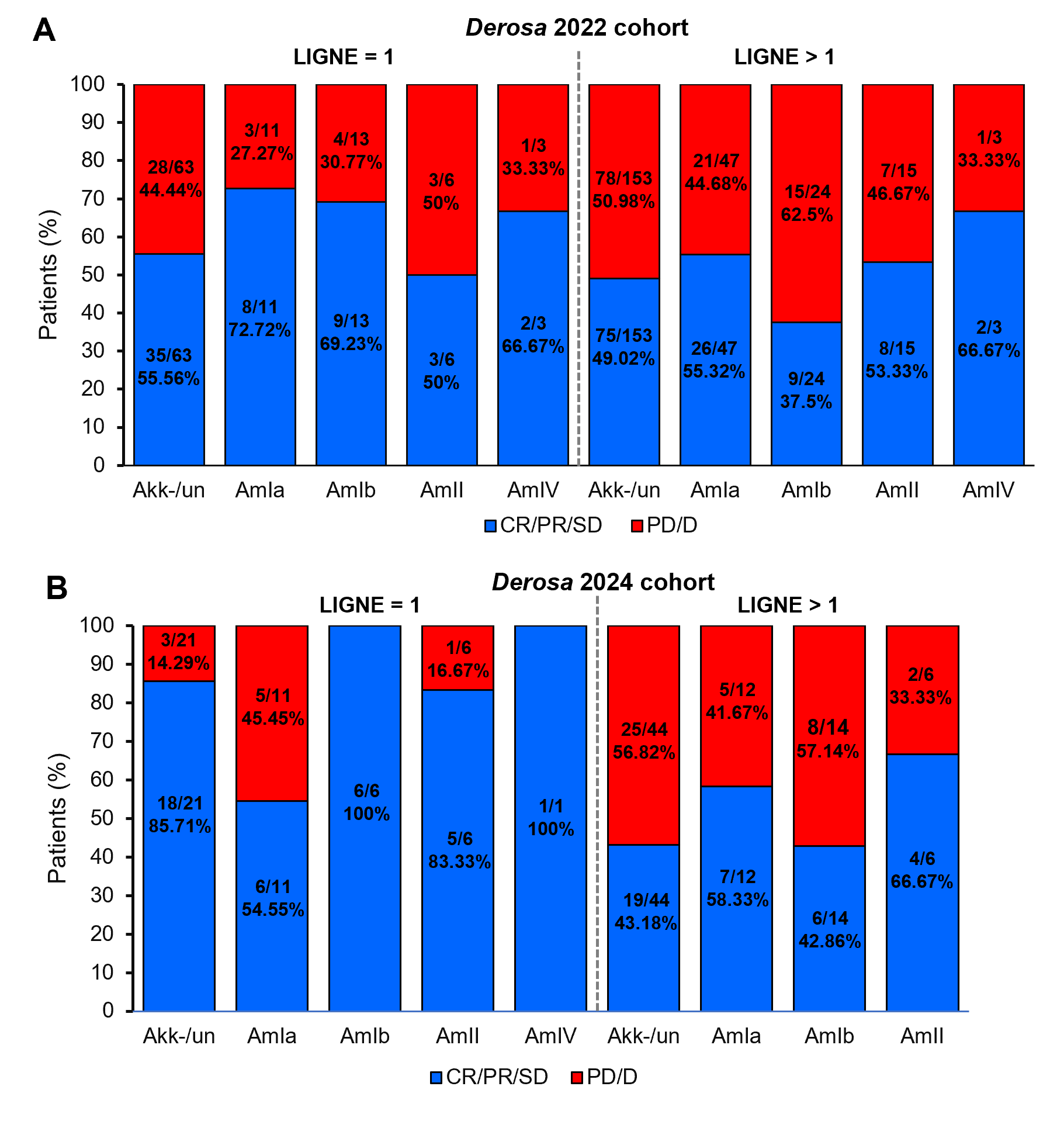

**Figure S6. The effect of LIGNE on the response to PD-1 blockade in patients carrying different Akk phylogroups in two NSCLC cohorts.** Differences in response to PD-1 blockade between patients carrying different Akk phylogroups among patients with LIGNE =1 or LINGE > 1 in *Derosa* 2022 cohort (C3) (**A**) and in *Derosa* 2024 cohort (C4) (**B**). Akk-/un represents for the samples that are either detected as Akk negative using MetaPhlAn 4 or detected as Akk positive in MetaPhIAN 4 but not in PanPhlan.

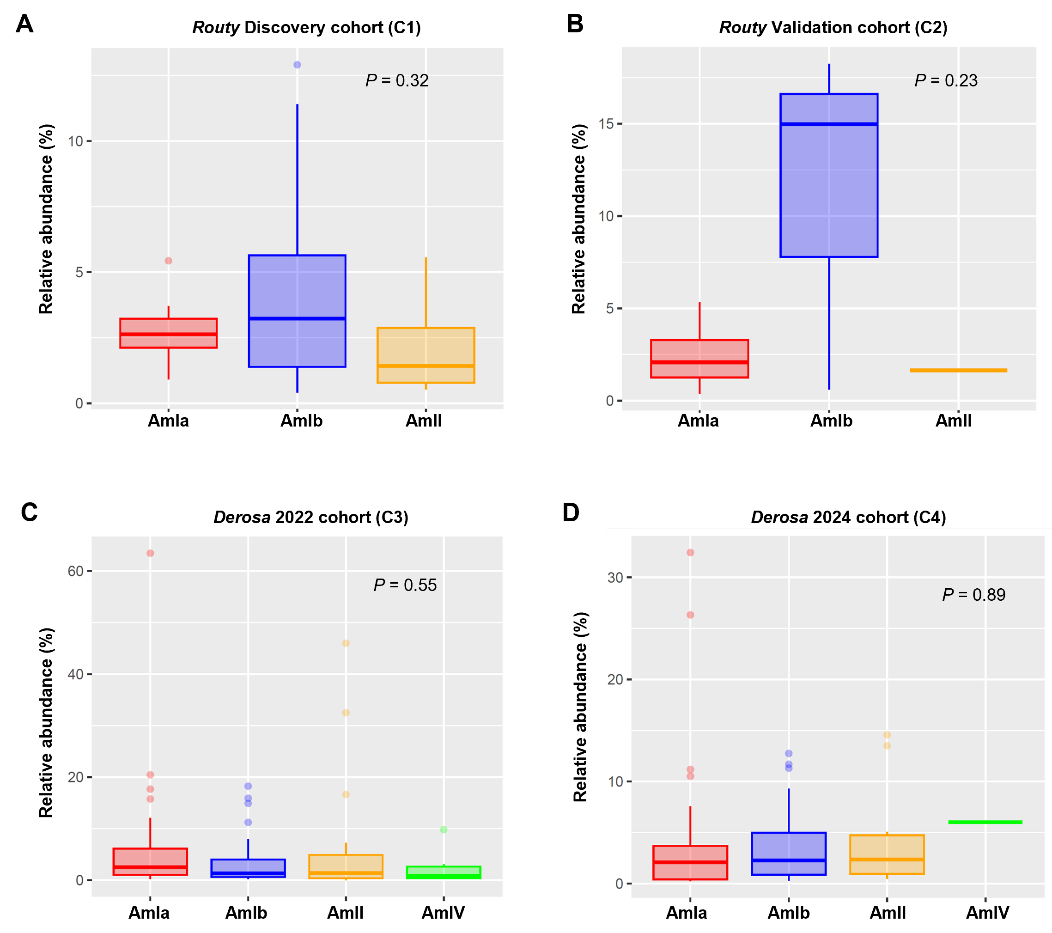

**Figure S7. Relative abundance of Akk phylogroups across four cohorts. There were no significant differences in relative abundance between Akk phylogroups in** *Routy* Discovery cohort (C1) (**A**), *Routy* Validation cohort (C2) (**B**), *Derosa* 2022 cohort (C3) (**C**), and *Derosa* 2024 cohort (C4) (**D**). The statistical analysis was conducted using the one-way ANOVA.

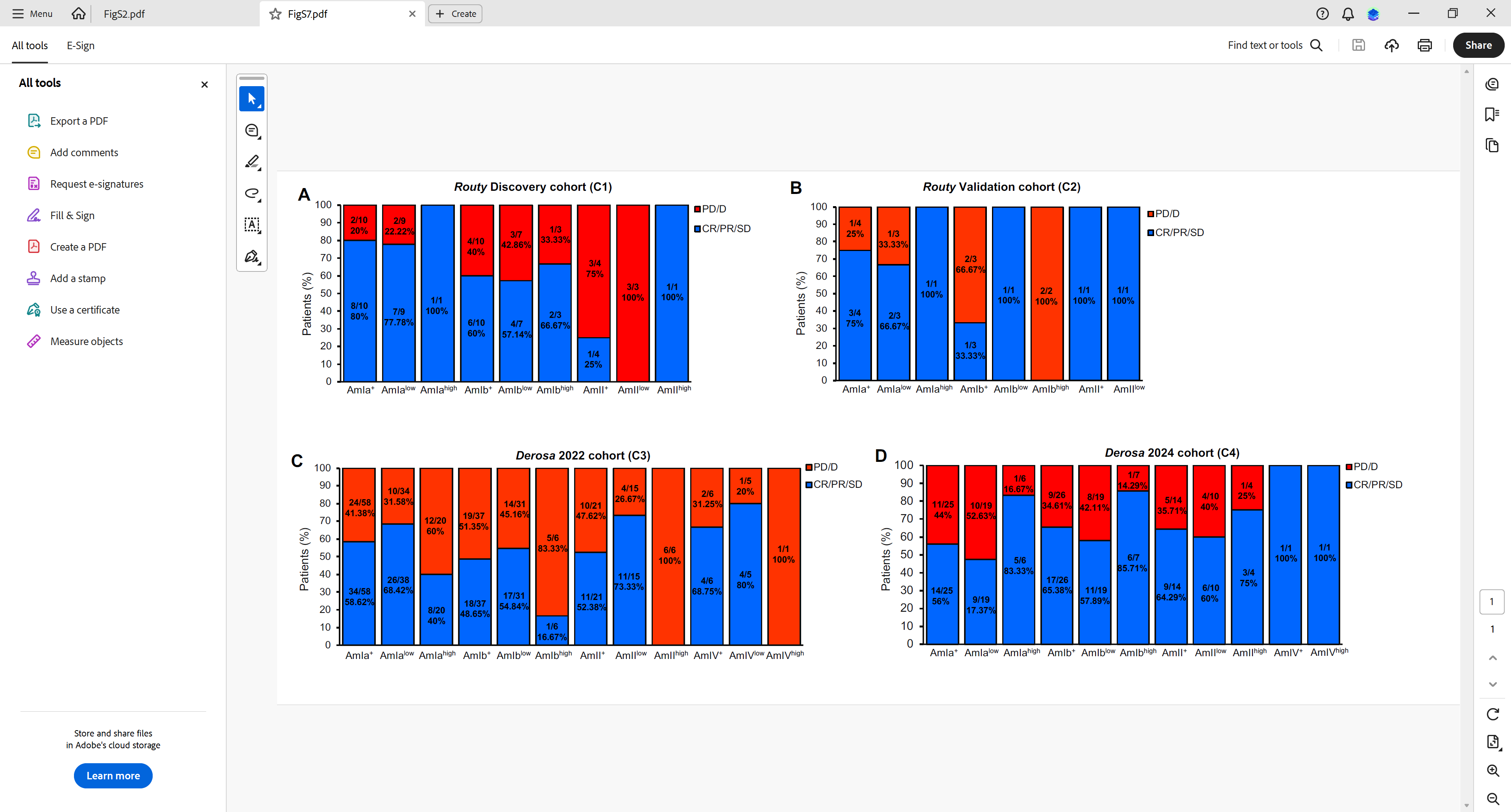

**Figure S8. NSCLC patients carrying different relative abundance of phylogroup of Akk varied in response to ICI treatment.** Differences in response to PD-1 blockade between patients carrying high (>4.799%) or low (< 4.799%) relative abundance of Akk phylogroups in *Routy* Discovery cohort (C1) (**A**), *Routy* Validation cohort (C2) (**B**), *Derosa* 2022 cohort (C3) (**C**), and *Derosa* 2024 cohort (C4) (**D**).

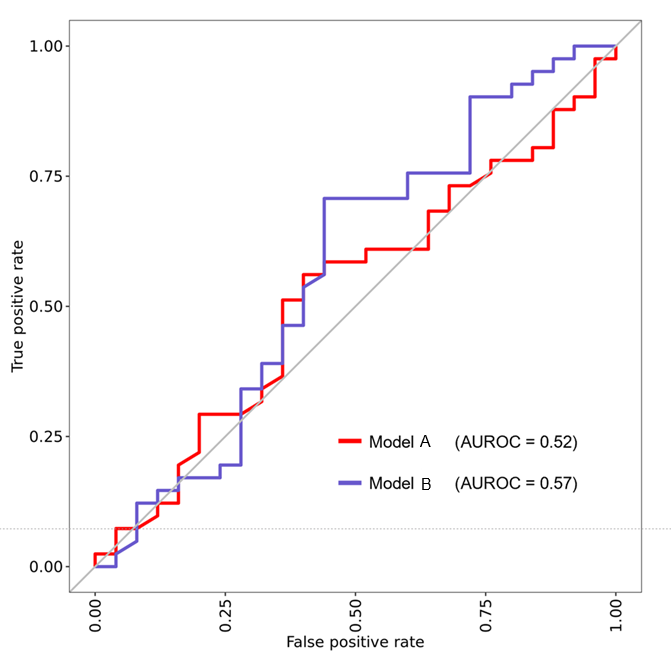

**Figure S9. Akk gene-based model improved performance to predict response to PD-1 blockade. (A) The area under the receiver operating characteristic (AUROC) of the random forest models trained to predict response using Akk phylogroup, patients’ metadata, number of clean reads, and without (model A) or with (model B) presence/absence of Akk genes in the combined C1 (n = 24), C2 cohort (n = 8), and C3 cohort (n = 122), and tested on C4 cohort LIGNE =1 patients (n = 66).**

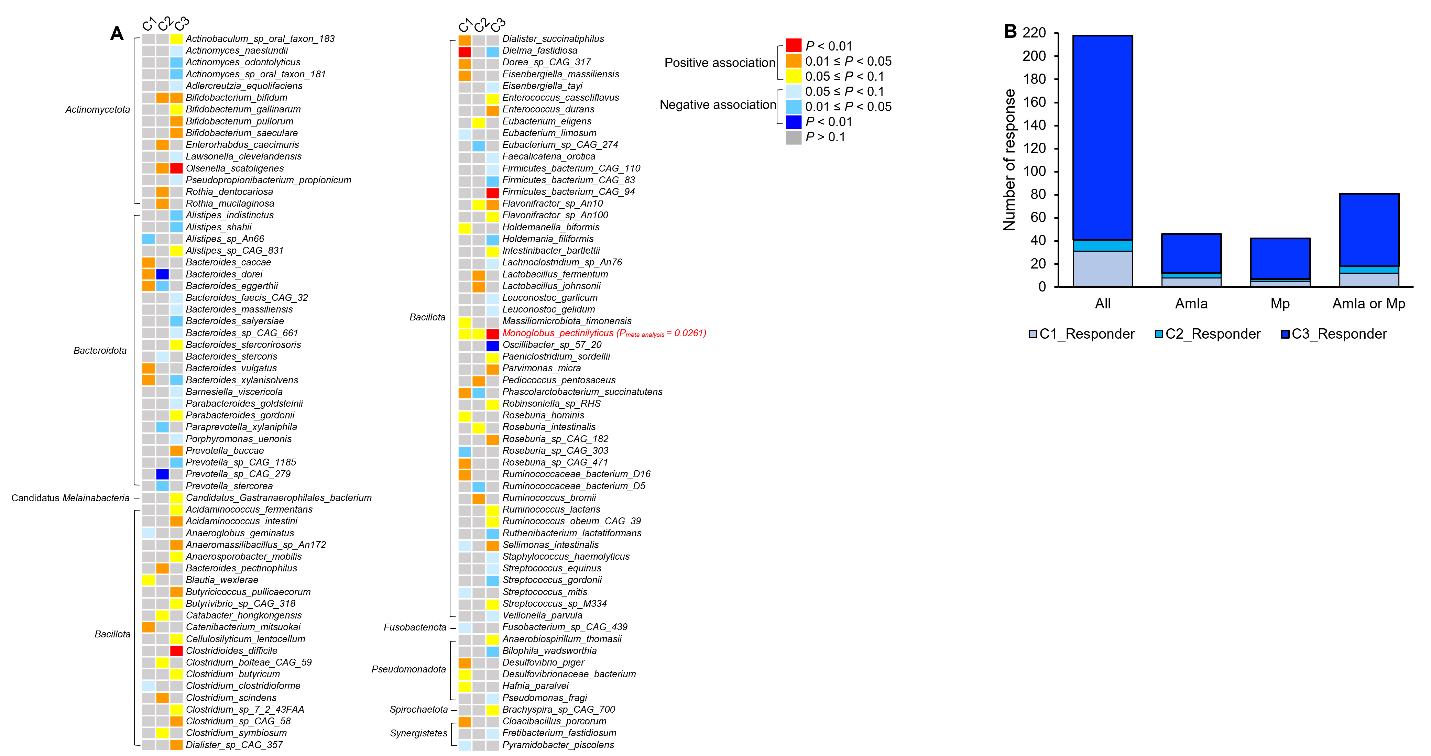

**Figure S10. The contribution of additional bacterial species to improving the predictive performance of response to PD-1 blockade.** (**A**) The heatmap showing the association between presence or absence of individual bacterial species and response to PD-1 blockade based on the generalized linear model. For each cohort, presence of individual bacterial species with sequencing depth and patients’ metadata were included as independent variables and response (Responders including CR, PR, and stable SD or Nonresponder including PD or D) was served as dependent variable. (**B**) The bargraph showing the increase of patients being predicted with the addition of *Monoglobus pectinilyticus* to AmIa in the model.

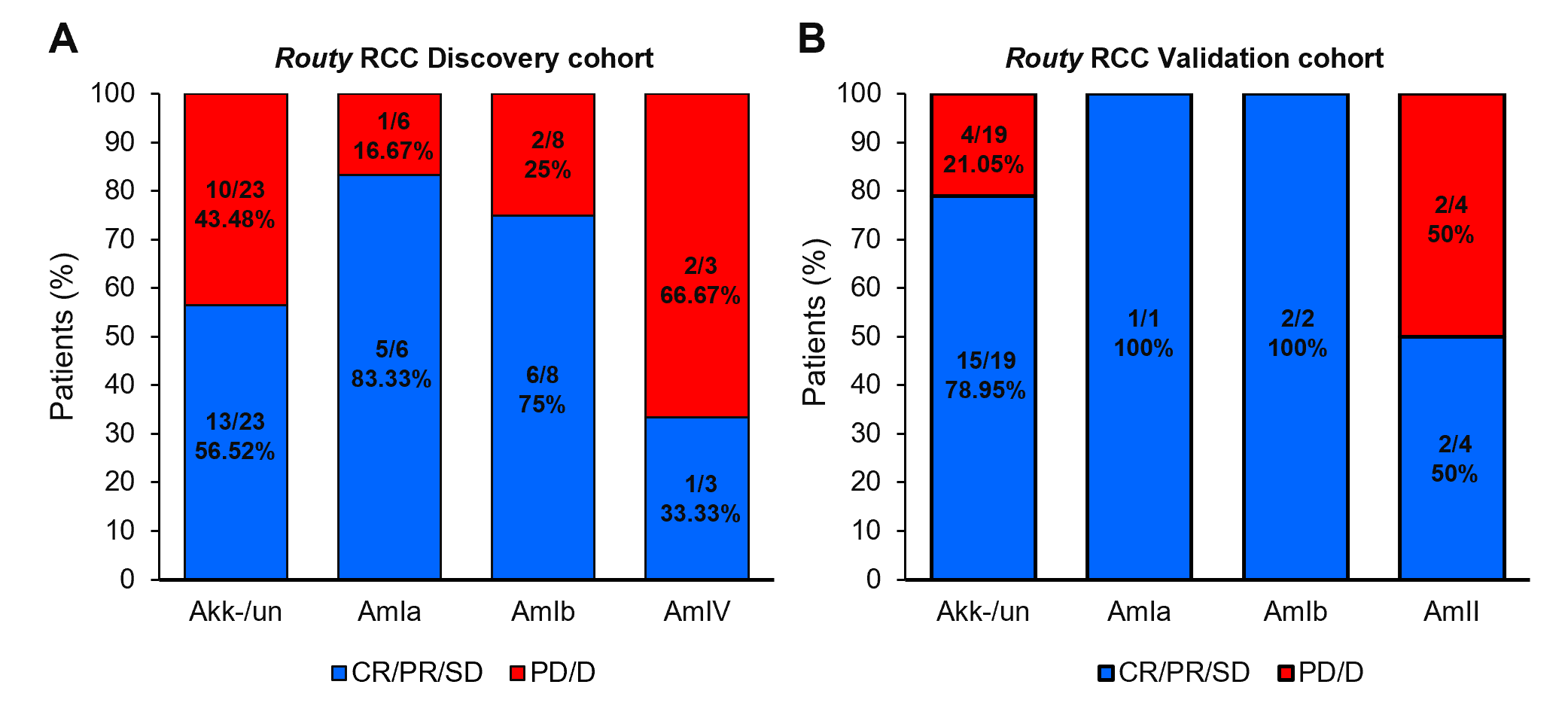

**Figure S11. Response to PD-1 blockade against renal cell carcinoma in patients carrying different phylogroups of Akk.** Differences in response to PD-1 blockade in RCC patients carrying different phylogroups of Akk in *Routy* RCC Discovery cohort (**A**) and *Routy* RCC Validation cohort (**B**). Akk-/un represents for the samples that are either detected as Akk negative using MetaPhlAn 4 or detected as Akk positive in MetaPhIAN 4 but not in PanPhlan.
